# Cyclophilin A protects HIV-1 from restriction by human TRIM5α

**DOI:** 10.1101/587907

**Authors:** Kyusik Kim, Ann Dauphin, Sevnur Komurlu, Leonid Yurkovetskiy, William E. Diehl, Sean M. McCauley, Claudia Carbone, Caterina Strambio-De-Castillia, Edward M. Campbell, Jeremy Luban

## Abstract

The capsid (CA) protein lattice of HIV-1 and other retroviruses encases viral genomic RNA and regulates steps that are essential to retroviral invasion of target cells, including reverse transcription, nuclear trafficking, and integration of viral cDNA into host chromosomal DNA^1^. Cyclophilin A (CypA), the first cellular protein reported to bind HIV-1 CA^2^, has interacted with invading lentiviruses related to HIV-1 for millions of years^3–7^. Disruption of the CA-CypA interaction decreases HIV-1 infectivity in human cells^8–12^, but stimulates infectivity in non-human primate cells^13–15^. Genetic and biochemical data suggest that CypA interaction with CA protects HIV-1 from a restriction factor in human cells^16–20^. Discovery of the CA-specific restriction factor TRIM5α^21^, and of TRIM5-CypA fusion genes that were independently generated at least four times in phylogeny^4,5,15,22–25^, pointed to human TRIM5α as the CypA-sensitive restriction factor. However, significant HIV-1 restriction by human TRIM5α^21^, let alone inhibition of such activity by CypA^26^, has not been detected. Here, exploiting reverse genetic tools optimized for primary human CD4^+^ T cells, macrophages, and dendritic cells, we demonstrate that disruption of the CA-CypA interaction renders HIV-1 susceptible to restriction by human TRIM5α, with the block occurring before reverse transcription. Identical findings were obtained with single-cycle vectors or with replication-competent HIV-1, including sexually-transmitted clones from sub-Saharan Africa. Endogenous TRIM5α was observed to associate with virion cores as they entered the macrophage cytoplasm, but only when the CA-CypA interaction was disrupted. These experiments resolve the long-standing mystery of the role of CypA in HIV-1 replication by demonstrating that this ubiquitous cellular protein shields HIV-1 from previously inapparent, but potent inhibition, imposed by human TRIM5α. Hopefully this reinvigorates development of CypA-inhibitors for treatment of HIV-1 and other CypA-dependent pathogens^27–30^.

---

To assess the role of TRIM5α and CypA in the primary human blood cell types that serve as targets for HIV-1 infection *in vivo*, lentiviral vectors were optimized for titer and knockdown efficiency in these cells^26,31-34^. Human macrophages, dendritic cells, and CD4^+^ T cells were transduced with lentivectors bearing a puromycin resistance cassette and shRNAs targeting either TRIM5 or luciferase (Luc) as a control. After three days of selection in puromycin, knockdown was confirmed by RT-qPCR for TRIM5 mRNA, and by rescue of N-MLV restriction (Extended Data Fig. 2a-c), as done previously^26,31^. TRIM5 and Luc control knockdown cells were then challenged with single-cycle, VSV G-pseudotyped, HIV-1-GFP reporter vectors. Three days later, the percentage of GFP+ cells was assessed by flow cytometry as a measure of infectivity (Extended Data Fig. 1 shows the gating strategy).

As compared with Luc control knockdown, TRIM5 knockdown had minimal effect on HIV-1 transduction efficiency in macrophages, dendritic cells, or CD4^+^ T cells (Fig. 1a-d; Extended Data Fig. 3a). Infectivity of HIV-1 CA-P90A, a mutant that disrupts CypA binding^8,9^, was significantly attenuated in control knockdown cells generated with all three cell types (Fig. 1a-d). The effect was evident in cells from all blood donors tested (at least three blood donors per condition) and over a 100-fold range in challenge vector titer (Extended Data Fig. 3a). TRIM5 knockdown in macrophages, dendritic cells, or CD4^+^ T cells increased CA-P90A infectivity (Fig. 1a-d; Extended Data Fig 3a). Results were the same whether challenge was with a three-plasmid vector system based on the clade B HIV-1_NL4-3_ lab strain^33,35^ (Fig. 1a-c), or a two plasmid vector system based on the clade C HIV-1_ZM249M_ transmission-founder strain from Zambia^33,36^ (Fig. 1d).

**Fig. 1.**
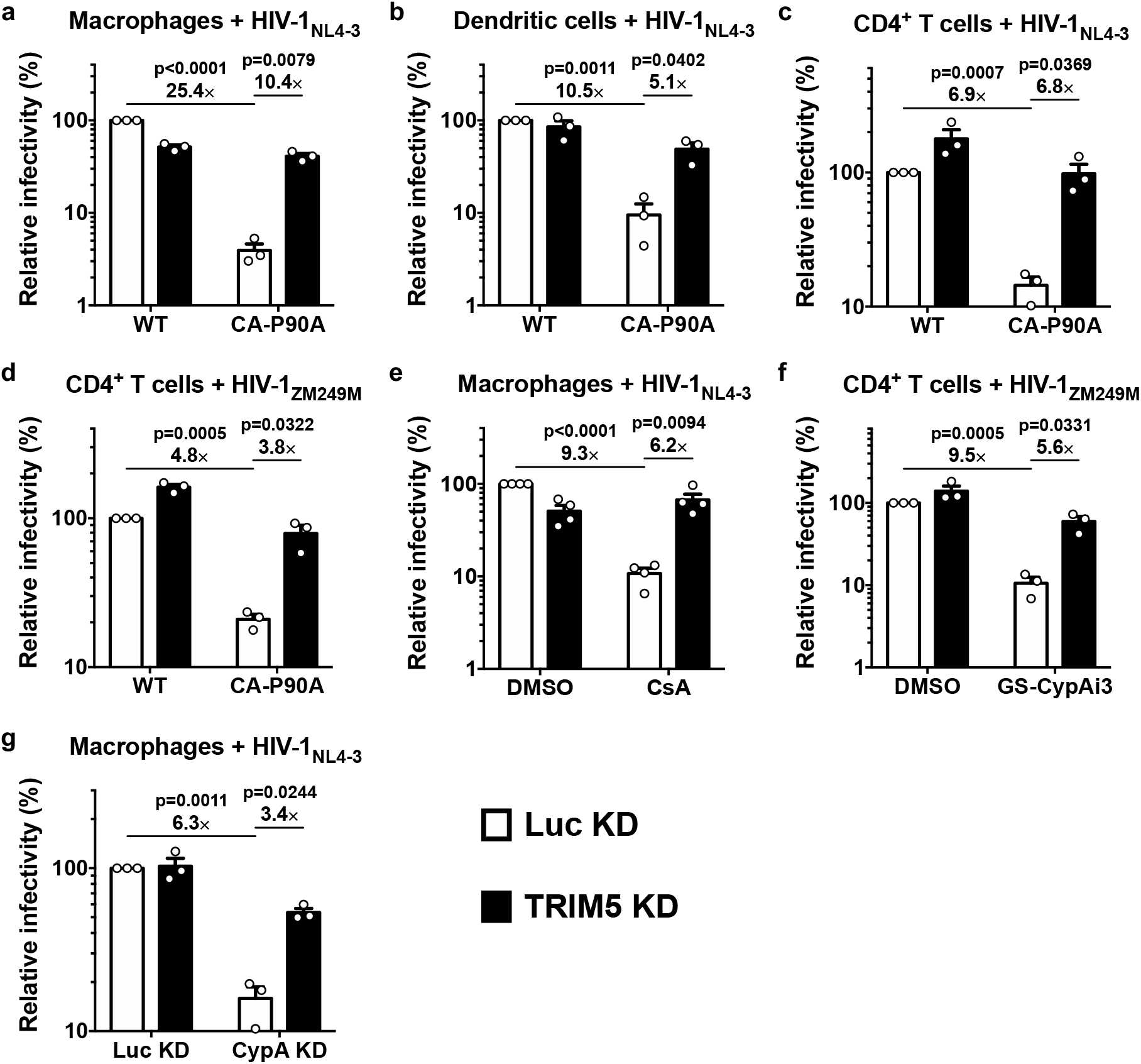
Disruption of the CA-CypA interaction in primary human blood cells renders HIV-1 susceptible to restriction by TRIM5. **a**, Macrophages, **b**, dendritic cells, or **c** and **d**, CD4^+^ T cells, were selected after transduction with a lentivirus expressing shRNA targeting TRIM5 or Luc control, and challenged with single-cycle, VSV G-pseudotyped, HIV-1_NL4-3_GFP (**a-c**) or HIV-1_ZM249M_GFP (**d**), bearing WT CA or CA-P90A (mean ± SEM, n = 3 donors for each), **e**, TRIM5 knockdown or Luc knockdown macrophages were challenged with HIV-1_NL4-3_GFP in the presence of 8 μM CsA or DMSO solvent (mean ± SEM, n = 4 donors), **f**, TRIM5 knockdown or Luc knockdown CD4^+^ T cells were challenged with HIV-1_ZM249M_GFP in the presence of 2.5 μM GS-CypAi3 or DMSO solvent (mean ± SEM, n = 3 donors), **g**, Macrophages were transduced simultaneously with two vectors expressing shRNAs, as indicated, and selected with puromycin and blasticidin. Cells were then challenged with HIV-1_NL4-3_GFP (mean ± SEM, n = 3 donors). The percentage of GFP^+^ cells was assessed by flow cytometry and normalized to WT in Luc control knockdown cells in all cases. Significance was determined by two-tailed, paired t-test.

Given previous reports that CypA and TRIM5 act independently to regulate HIV-1 transduction of immortalized cell lines^26^, the rescue of CA-P90A infectivity by TRIM5 knockdown in primary human blood cells was surprising. Complementary pharmacologic and reverse genetic approaches were therefore taken to disrupt the CA-CypA interaction. For pharmacologic disruption, cells were incubated in media containing small molecules that compete with CA for binding to CypA^2,8–10,37^. As compared with DMSO solvent alone, cyclosporine A (CsA) reduced HIV-1 transduction efficiency in control Luc knockdown macrophages (Fig. 1e). In contrast, cyclosporine H, an analogue with 1,000-fold lower affinity for CypA^38^, caused slight increase in HIV-1 infection (Extended Data Fig. 3b). Either of two non-immunosuppressive CypA inhibitors derived from sanglifehrin A^37^, decreased HIV-1 transduction efficiency in primary CD4^+^ T cells (Fig. 1f; Extended Data Fig. 3c-e). TRIM5 knockdown reversed the HIV-1 inhibition in macrophages caused by CsA (Fig. 1e), or in CD4^+^ T cells caused by the sanglifehrin A-derivatives (Fig. 1f; Extended Data Fig. 3c-e).

To disrupt the CA-CypA interaction using a genetic approach, macrophages were transduced with two vectors. The first vector conferred puromycin resistance and expressed shRNAs targeting either TRIM5 or Luc. The second vector conferred blasticidin resistance and expressed shRNAs targeting either CypA or Luc. After simultaneous transduction with pairs of these vectors, macrophages were selected for three days in both puromycin and blasticidin and then challenged with single-cycle, VSV G-pseudotyped, HIV-1-GFP reporter vector bearing wild-type CA. As compared with the control Luc knockdown, CypA knockdown reduced CypA protein levels ~70% (Extended Data Fig. 2d). Like CA-P90A and the small molecule inhibitors, CypA knockdown decreased transduction efficiency (Fig. 1g). This effect was rescued by simultaneous knockdown of TRIM5 (Fig. 1e) without restoring CypA protein levels (Extended Data Fig. 2d).

Though the shRNA that targets TRIM5 is distinguished from the next most similar sequence in the human genome (GRCh38) by multiple mismatches, off-target effects are theoretically possible. To test whether TRIM5 is sufficient to explain the HIV-1 restriction activity associated with CA-CypA disruption, a vector was designed based on the ubiquitin fusion technique^39^, that expresses a tripartite fusion of puromycin N-acetyl transferase (PAC), ubiquitin^K48R^, and coding sequence for a protein of interest, in addition to an shRNA (Fig. 2a). Four variants of the plasmid were engineered in which the shRNA targeted either TRIM5 or Luc, with or without a TRIM5α coding sequence that bears mismatches in the shRNA target sequence (Fig. 2a).

**Fig. 2.**
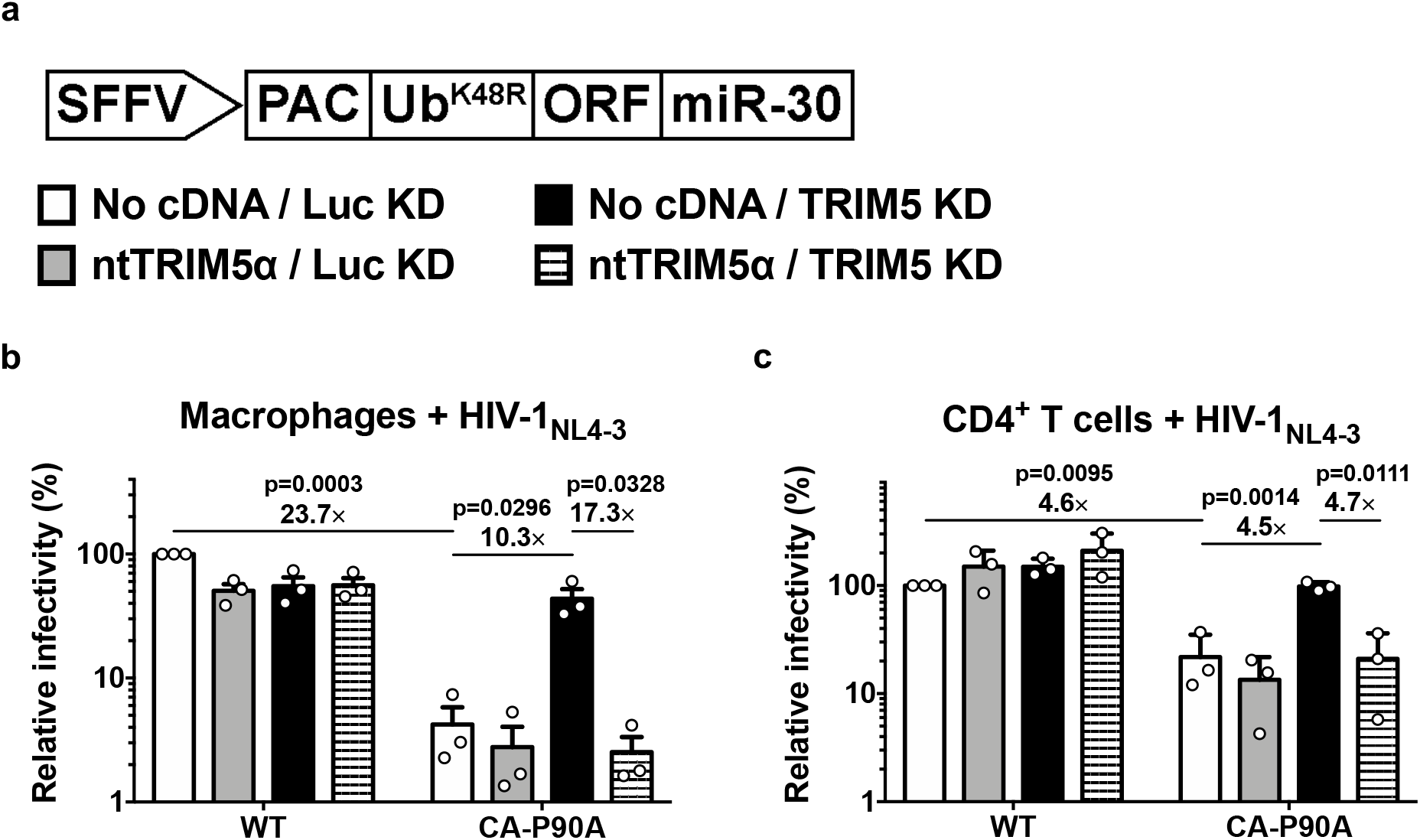
Human TRIM5α is sufficient to explain the HIV-1 inhibition that results from disruption of the CA-CypA interaction. **a**, Schematic representation of the all-in-one shRNA-rescue lentivector, in which the SFFV promoter expresses a tripartite fusion of puromycin N-acetyl transferase (PAC), the K48R mutant of ubiquitin, and an open reading frame for a gene of interest (ORF), as well as a miR-30 based shRNA (miR-30). **b** and **c**, All-in-one lentivectors encoding the shRNAs and ORFs indicated in **a** were used to transduce macrophages (**b**) or CD4^+^ T cells (**c**). The percentage of GFP-expressing cells was measured by flow cytometry and normalized to the values for No cDNA/Luc KD cells challenged with WT CA; mean ± SEM, n = 3 donors for each. Significance was determined by two-tailed, paired t-test.

Macrophages (Fig. 2b) and CD4^+^ T cells (Fig. 2c) were transduced with each of the four variants of the shRNA tripartite fusion vector and selected for three days with puromycin. Cells were then challenged with the three-plasmid HIV-1-GFP reporter vector used in Fig. 1a, and assessed by flow cytometry for percent GFP positive cells three days later. As in Fig. 1, the infectivity of vector bearing wild-type CA was minimally affected by TRIM5 knockdown, or by TRIM5 overexpression (Fig. 2b, c). As compared to the wild-type, the infectivity of vector bearing CA-P90A was decreased (Fig. 2b, c), and the infectivity of this mutant was rescued by TRIM5 shRNA (Fig. 2b, c). In the presence of the shRNA targeting TRIM5, delivery of TRIM5α coding sequence bearing shRNA target-site mismatches restored restriction activity to the control level (Fig. 2b, c). These results demonstrate that, in primary human blood cells, human TRIM5α is sufficient to restrict HIV-1 transduction, but only when the CA-CypA interaction is disrupted.

To determine at which step in the virus life cycle human TRIM5α inhibits HIV-1 when the CA-CypA interaction is disrupted, HIV-1 cDNA resulting from reverse transcription was assessed by qPCR. Macrophages and CD4^+^ T cells were stably transduced and selected with vector expressing shRNAs targeting TRIM5 or Luc control (Fig 3a, b), or with each of the four variants of the shRNA tripartite fusion vector (Fig. 3c, d). Cells were then challenged with an HIV-1 reporter vector that has the 34 basepair loxP sequence in U3 to distinguish reporter vector transcripts from those of the shRNA lentivector^40^. DNA was collected 20 hrs post-challenge and qPCR was performed using primers specific for full-length linear HIV-1 cDNA (late RT). In all experiments, reporter vector bearing the RT-D185K/D186L loss-of-function mutation^40^ was included as a control for background signal not due to nascent reverse transcription (Fig. 3).

**Fig. 3.**
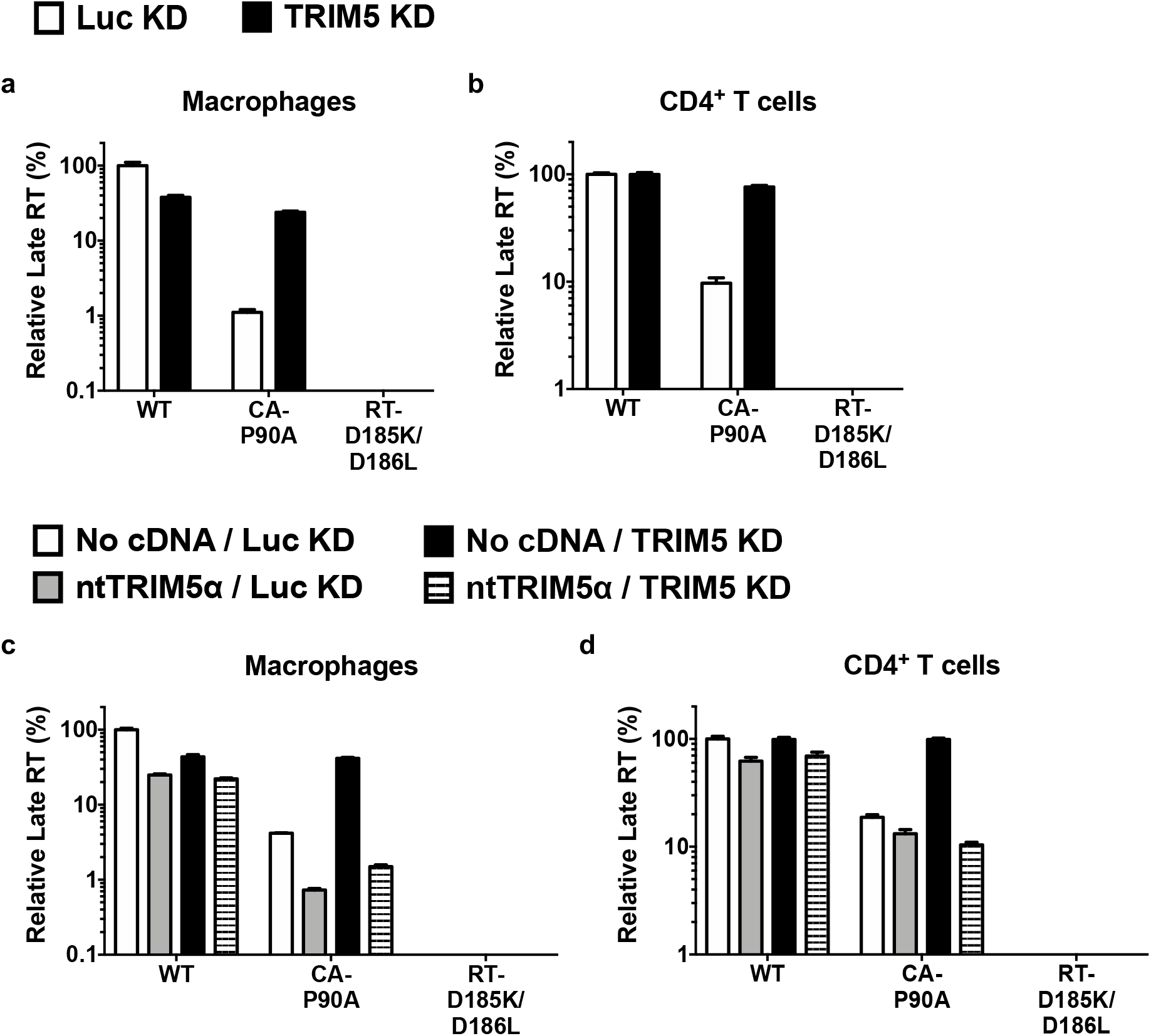
CypA protects HIV-1 from restriction by human TRIM5α prior to completion of reverse transcription. **a-d**, TRIM5 knockdown or Luc knockdown macrophages (**a**) or CD4^+^ T cells (**b**), or macrophages (**c**) or CD4^+^ T cells (**d**) transduced with the all-in-one shRNA-rescue lentivectors described in Fig. 2 were challenged with HIV-1_NL4-3_GFP containing WT CA or CA-P90A, as indicated. DNA was extracted 20 hrs post-challenge and late products of reverse transcription (RT) were assessed by qPCR (mean of triplicates ± SEM). RT-D185K/D186L mutant virus was used as a control for background.

In Luc control knockdown macrophages and CD4^+^ T cells, viral cDNA was reduced by CA-P90A, and this reduction was restored by TRIM5 knockdown (Fig. 3a, b). Viral cDNA was also reduced by CA-P90A in either cell type transduced with the control shRNA tripartite fusion vector (Fig. 3c, d). TRIM5 shRNA rescued the cDNA (Fig. 3a-d), and rescue of TRIM5α with the non-targetable coding sequence again decreased the CA-P90A cDNA (Fig. 3c, d). These results demonstrate that, when the CA-CypA interaction is disrupted, human TRIM5α blocks HIV-1 at an early step of viral infection, prior to completion of reverse transcription.

To determine if TRIM5α associates with HIV-1 CA in cells when the CA-CypA interaction is disrupted, primary human macrophages were stably transduced with TRIM5 shRNA or Luc shRNA and then challenged for 2 hrs with wild-type HIV-1 reporter vector, in the presence or absence of CsA. Cells were fixed and subjected to the proximity ligation assay (PLA) with antibodies specific for HIV-1 CA and endogenous human TRIM5α. When cells were challenged with HIV-1 in the absence of CsA, very few puncta were detected (Fig. 4a, b; Extended Data Fig. 4). Similarly, few puncta were detected when cells were treated with CsA in the absence of HIV-1 challenge. In contrast, when cells were challenged with HIV-1 in the presence of CsA, multiple puncta were detected (Fig. 4a, b), an increase of at least 20-fold in the average number of puncta per cell over the background (Fig. 4b; Extended Data Fig. 4). TRIM5 knockdown eliminated the puncta (Fig. 4a, b; Extended Data Fig. 4), indicating that the PLA signal was dependent upon TRIM5 expression. These results indicate that, in acutely infected primary human macrophages, endogenous human TRIM5α associates with HIV-1 CA when the CA-CypA interaction is disrupted.

**Fig. 4.**
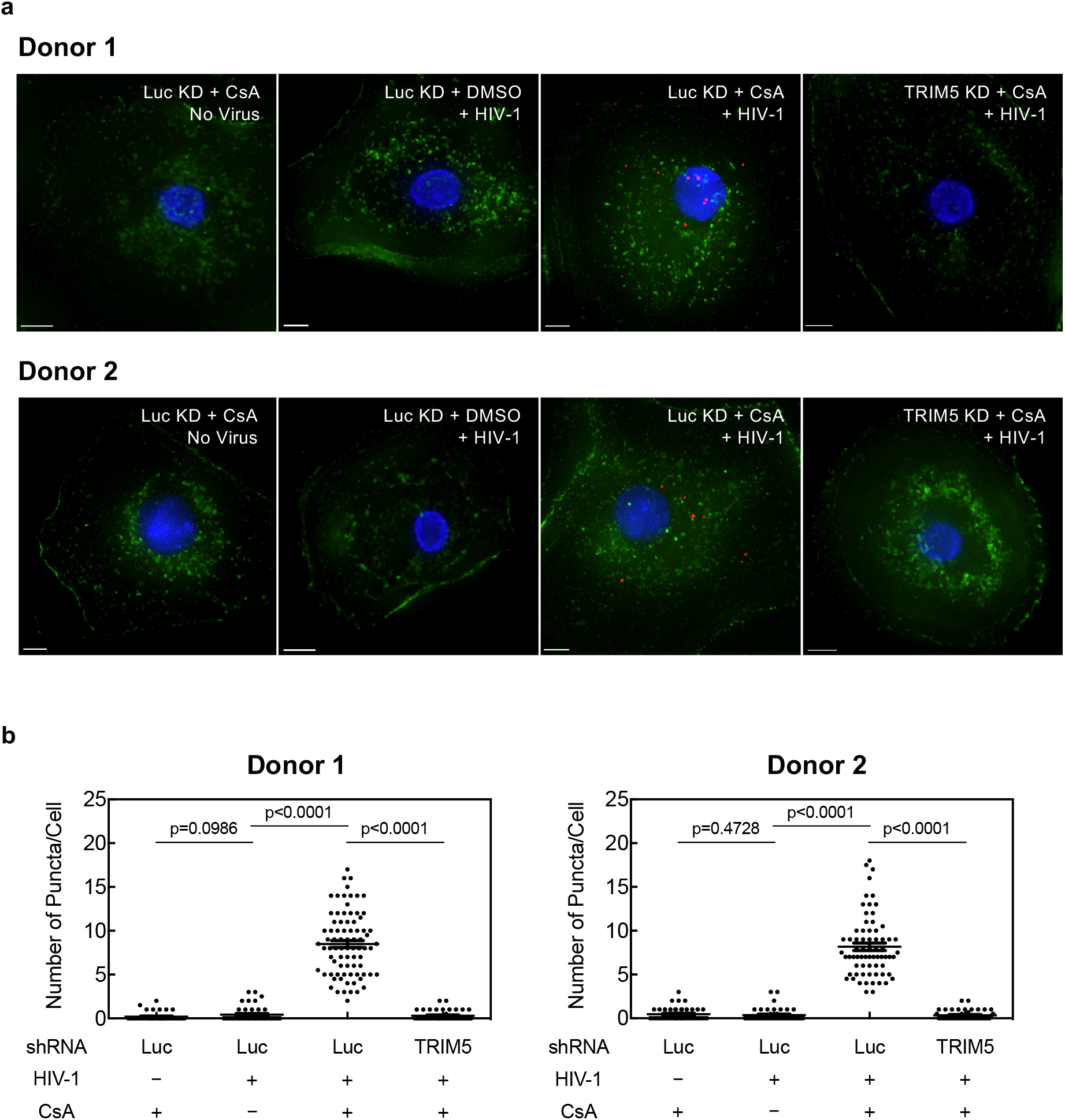
Endogenous TRIM5α in primary human macrophages associates with HIV-1 CA after acute challenge but only when the CA-CypA interaction is disrupted. **a**, Macrophages from two blood donors were transduced and selected with vector bearing shRNA targeting either TRIM5 or Luc. Cells were then challenged for 2 hrs with VSV G-pseudotyped, HIV-1_NL4-3_GFP, in the presence of 5 μM CsA or DMSO solvent. Cells were fixed and the PLA assay was performed with antibodies against HIV-1 CA and TRIM5α. Representative images show PLA puncta (red), nuclei stained with Hoechst (blue), and actin filaments stained with phalloidin (green). **b**, The number of puncta per cell in the PLA assay, after analysis of at least 45 cells, per condition, in triplicate (mean ± SEM). Significance was determined by two-tailed, unpaired t-test.

The above experiments used exogenous, single-cycle HIV-1 vectors. The effect of CypA on HIV-1 restriction by human TRIM5α was therefore evaluated next using replication-competent HIV-1 in a context where the virus spreads cell-to-cell. Primary human macrophages were challenged with clade B HIV-1 bearing a macrophage-tropic *env* (HIV-1_MAC_) and replication was monitored for 14 days by measuring the accumulation of reverse transcriptase activity in the supernatant. As in the single cycle experiments (Fig. 1), TRIM5 knockdown itself had little effect on HIV-1 replication (Fig. 5a, b). Disruption of the CA-CypA interaction with CsA (Fig. 5a) or with shRNA targeting CypA (Fig. 5b) effectively suppressed viral spread in the culture, and, in both cases, replication kinetics was completely restored to the control level by shRNA targeting TRIM5 (Fig. 5a, b). Primary CD4^+^ T cells were then challenged with a clade C transmission/founder virus (HIV-1_ZM249M_). As observed in macrophages, TRIM5 knockdown alone had minimal effect on wild-type HIV-1 replication (Fig. 5c, d). No viral replication was detectable when the CA-CypA interaction was disrupted by the small molecule GS-CypAi3 (Fig. 5c) or by the presence of CA-P90A in HIV-1 (Fig. 5d); in both cases shRNA targeting TRIM5 rescued replication kinetics to the level of controls (Fig. 5c, d).

**Fig. 5.**
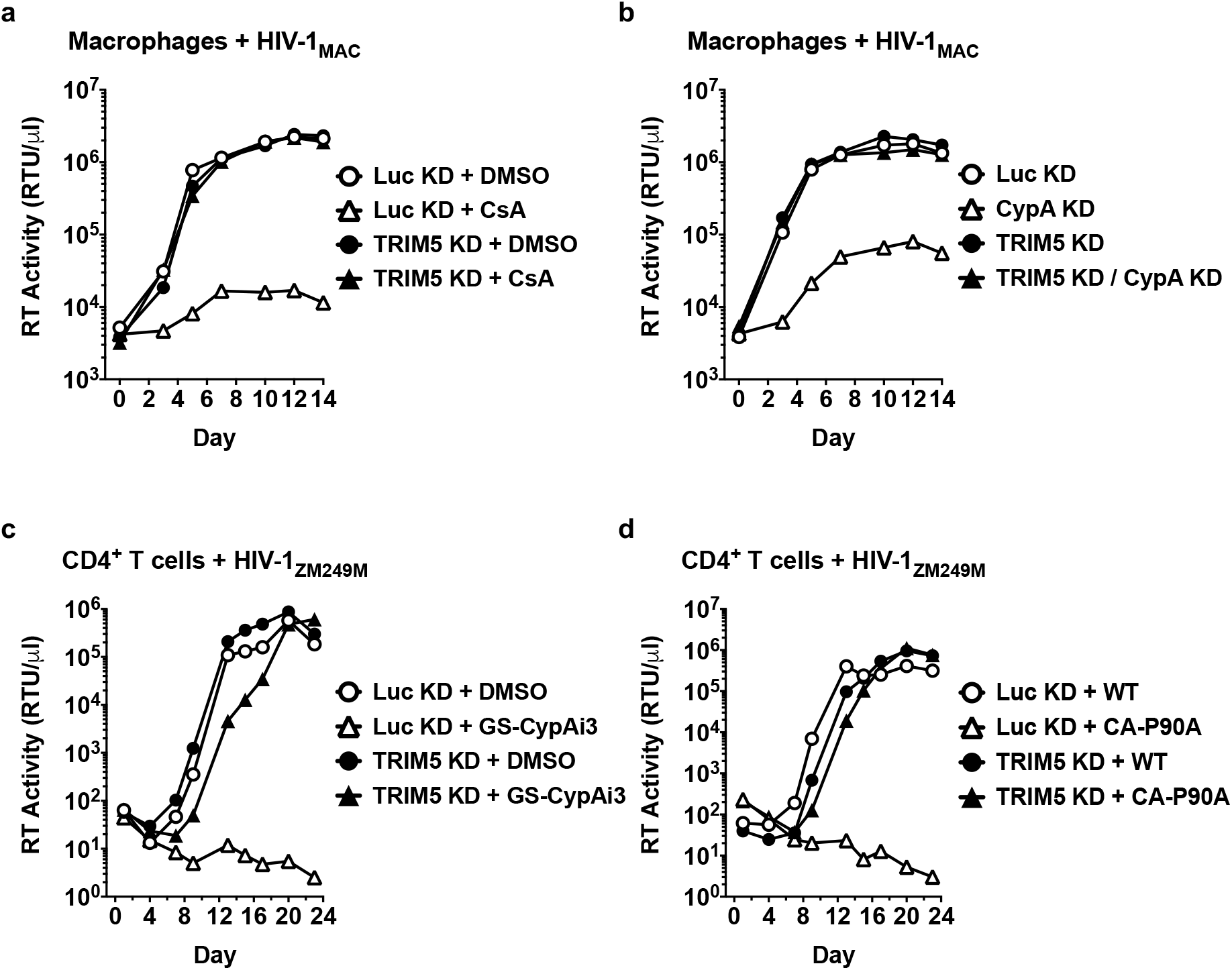
Endogenous TRIM5α suppresses spreading infection of HIV-1 in human primary macrophages and CD4^+^ T cells when the CA-CypA interaction is disrupted. **a** and **b**, Spreading infection of HIV-1_MAC_ in TRIM5 or Luc knockdown macrophages with 5 μM CsA (**a**) or with vectors bearing shRNAs targeting CypA or Luc (**b**), as indicated. **c** and **d**, Spreading infection of HIV-1_ZM249M_ in CD4^+^ T cells expressing shRNA targeting TRIM5 or Luc with 2.5 μM GS-CypAi3 (**c**) or when challenged with virus bearing CA-P90A (**d**), as indicated. HIV-1 replication was monitored by measuring reverse transcriptase (RT) activity in the culture supernatant over time. Data shown are representative of experiments in two blood donor cells for each condition.

The experiments presented here demonstrate that, in primary human blood cells, HIV-1 exploits CypA to evade CA recognition by, and the antiviral activity of, endogenous TRIM5α. This answers the long-standing question of how CypA promotes HIV-1 infection and clearly establishes that, in the absence of CypA, human TRIM5α potently restricts HIV-1. Conservation of the lentiviral CA-CypA interaction across millions of years of evolution is likely a result of selective pressure applied by TRIM5α orthologues encoded by host species that are otherwise permissive for lentiviral replication. Finally, the results here, in which primary human blood cells were challenged by clones of HIV-1 that were isolated after human-to-human transmission, indicate that, by rendering HIV-1 susceptible to the potent antiviral activity of TRIM5α, non-immunosuppressive CypA inhibitors have potential to make an important contribution to anti-HIV-1 drug cocktails.

## Methods

### Plasmids

All plasmids used here are described in Supplementary Table 1, and are available, along with full sequences, at https://www.addgene.org/Jeremy_Luban/.

### Human blood

Leukopaks were obtained from anonymous, healthy, blood donors (New York Biologics, Southhampton, NY). These experiments were reviewed by the University of Massachusetts Medical School Institutional Review Board, and declared non-human subjects research, according to NIH guidelines (http://grants.nih.gov/grants/policy/hs/faqs_aps_definitions.htm).

### Cell culture

All cells were cultured in humidified, 5% CO_2_ incubators at 37°C. HEK293 cells (ATCC) were cultured in DMEM supplemented with 10% heat-inactivated FBS, 1 mM sodium pyruvate, 20 mM GlutaMAX^™^-I, 1× MEM non-essential amino acids, and 25 mM HEPES, pH 7.2 (DMEM-FBS complete). PBMCs were isolated from leukopaks by gradient centrifugation on Lymphoprep (Axis-Shield PoC As, Oslo, Norway, catalogue #AXS-1114546). To generate dendritic cells (DCs) or macrophages, CD14+ mononuclear cells were enriched by positive selection using anti-CD14 antibody microbeads (Miltenyi, San Diego, CA, catalogue #130-050-201). Enriched CD14+ cells were plated in RPMI-1640, supplemented with 5% heat-inactivated human AB+ serum (Omega Scientific, Tarzana, CA), 1 mM sodium pyruvate, 20 mM GlutaMAX^™^-I, 1× MEM non-essential amino acids, and 25 mM HEPES pH 7.2 (RPMI-HS complete), at a density of 10^6^ cells/mL for macrophages or 2×10^6^ cells/mL for DCs. To differentiate CD14+ cells into macrophages, 1:100 GM-CSF-conditioned media was added. To differentiate CD14+ cells into DCs, 1:100 cytokine-conditioned media containing human GM-CSF and human IL-4 was added. GM-CSF and IL-4 were produced from HEK293 cells transduced with pAIP-hGMCSF-co (Addgene #74168) or pAIP-hIL4-co (Addgene #74169), as previously described^31,33^. CD4^+^ T cells were isolated from CD14-depleted PBMCs using anti-CD4 antibody microbeads (Miltenyi, catalogue #130-045-101); enrichment was typically >90%, as assessed by measuring the percentage of CD3^+^/CD4^+^ cells via flow cytometry with FITC-anti-CD3 (Biolegend, San Diego, CA, catalogue #317306) and APC-anti-CD4 (Biolegend, catalogue #317416). The cells were cultured in RPMI-1640 supplemented with 10% heat-inactivated FBS, 1 mM sodium pyruvate, 20 mM GlutaMAX^™^-I, 1× MEM non-essential amino acids, and 25 mM HEPES pH 7.2 (RPMI-FBS complete) with 50 U/mL hIL-2 (NIH AIDS Reagent Program, catalogue #136).

### Virus production

24 hrs prior to transfection, 6 × 10^5^ HEK293 cells were plated per well in 6-well plates. All transfections used 2.49 μg plasmid DNA with 6.25 μL TransIT LT1 transfection reagent (Mirus, Madison, WI), in 250 μL Opti-MEM (Gibco). 2.49 μg of replication-competent HIV-1 provirus DNA was transfected. For two-part, single-cycle vector, 2.18 μg of env-defective HIV-1 provirus was co-transfected with 0.31 μg pMD2.G VSV G plasmid. Three-part, single-cycle vectors were produced by co-transfecting 1.25 μg minimal lentivector genome plasmid (either pALPS-GFP, pWPTS-GFP, pLXIN-GFP, pAPM-CoE-D4-miR30, pABM-CoE-D4-miR30, or pPU-ORF-miR30), 0.93 μg *gag-pol* plasmid (either psPAX2, p8.9 NΔSB, pCIG3-N or pCIG3-B), and 0.31 μg pMD2.G VSV G plasmid. Vpx-containing SIV-VLPs were produced by transfection of 2.18 μg pSIV3+ and 0.31 μg pMD2.G plasmid. 16 hrs post-transfection, culture media was changed to the media specific for the cells to be transduced. Viral supernatant was harvested at 72 hrs, passed through a 0.45 μm filter, and stored at −80°C.

### Exogenous reverse transcriptase assay

5 μL transfection supernatant was mixed with 5 μL 0.25% Triton X-100, 50 mM KCl, 100 mM Tris-HCl pH 7.4, and 0.4 U/μL RiboLock RNase inhibitor, and then diluted 1:100 in 5 mM (NH_4_)_2_SO_4_, 20 mM KCl, and 20 mM Tris-HCl pH 8.3. 10 μL of this was then added to a single-step, RT-PCR assay with 35 nM MS2 RNA (IDT) as template, 500 nM of each primer (5’-TCCTGCTCAACTTCCTGTCGAG-3’ and 5’-CACAGGTCAAACCTCCTAGGAATG-3’), and 0.1 μL hot-start Taq DNA polymerase (Promega, Madison, WI) in 20 mM Tris-Cl pH 8.3, 5 mM (NH_4_)_2_SO_4_, 20 mM KCl, 5 mM MgCl_2_, 0.1 mg/ml BSA, 1/20,000 SYBR Green I (Invitrogen), and 200 μM dNTPs in total 20 μL reaction. The RT-PCR reaction was carried out in a Biorad CFX96 real-time PCR detection system with the following parameters: 42°C for 20 min, 95°C for 2 min, and 40 cycles [95°C for 5 sec, 60°C for 5 sec, 72°C for 15 sec, and acquisition at 80°C for 5 sec].

### Transduction with lentiviral knockdown vectors

For DCs, 2 × 10^6^ CD14+ monocytes/mL were transduced with 1:4 volume of SIV-VLPs and 1:4 volume of knockdown lentivector. For macrophages, 10^6^ CD14+ monocytes/mL were transduced with 1:8 volume of SIV-VLPs and 1:8 volume of knockdown lentivector. The Vpx-containing SIV-VLPs were added to these cultures to overcome a SAMHD1 block to lentiviral transduction^41,42^. Transduced cells were selected with 3 μg/mL puromycin (InvivoGen, San Diego, CA, catalogue #ant-pr-1), 10 μg/mL blasticidin (InvivoGen, catalogue #ant-bl-1), or both, for 3 days, starting 3 days post-transduction.

Following isolation with magnetic beads, human CD4^+^ T cells were cultured at 2 to 3 × 10^6^ cells/mL in RPMI-FBS complete, supplemented with 50 U/mL hIL-2, and stimulated with 5 μg/mL PHA-P (Sigma-Aldrich, catalogue #L-1668). Alternatively, CD4^+^ T cells at 10^6^ cells/mL were stimulated with 25 μL/mL ImmunoCult^™^ Human CD3/CD28 T Cell Activator (STEMCELL Technologies, Vancouver, Canada, catalogue #10991). At day 3 post-stimulation, T cells were replated at 2 to 3 × 10^6^ cells/mL in RPMI-FBS complete, with 50 U/mL hIL-2. Cells were transduced with 10^8^ RT units of viral vector per 10^6^ cells for 3 days, followed by selection with 2 μg/mL puromycin. After selection for 3 days, cells were re-stimulated with PHA-P, or with ImmunoCult^™^ Human CD3/CD28 T Cell Activator, for 3 days. The stimulated cells were then replated at 2 to 3 × 10^6^ cells/mL (PHA) or at 10^6^ cells/mL (CD3/CD28) in RPMI-FBS complete with 50 U/mL hIL-2, and challenged with lentiviral vectors for assessment of single-cycle infectivity or spreading infection. Fresh media containing hIL-2 was replenished every 2 to 3 days.

### Infectivity assay using single-cycle viruses

For human DCs, 2.5 × 10^5^ cells were seeded per well, in a 48-well plate, on the day of virus challenge. Media containing VSV G-pseudotyped lentiviral vector expressing GFP (HIV-1-GFP) was added to challenge cells in a total volume of 250 μL. For human macrophages, 2.5 × 10^5^ cells were seeded per well in a 24-well plate, and challenged with HIV-1-GFP in a total volume of 500 μL. 1:50 volume of SIV VLPs was also added to the medium during virus challenge of DCs or macrophages. To challenge human CD4^+^ cells activated with PHA, 5 × 10^5^ cells were plated per well in 96-well plate at 3 days after the second PHA stimulation. For CD4^+^ cells stimulated with CD3/CD28 activator, 2 × 10^5^ cells were plated in each well of a 96-well plate, 3 days after secondary stimulation. Cells were then challenged with GFP reporter viruses in a total volume of 200 μL. For all three cell types, 4 dilutions of viral stocks, from 10^5^ to 10^8^ RT units/mL, were used to challenge cells. Where indicated, cells were pre-treated with 8 μM cyclosporine A (CsA), 8 μM cyclosporine H (CsH), or 2.5 μM of non-immunosuppressive CypA inhibitors from Gilead (GS-CypAi3 or GS-CypAi48)^37^, for 1 hr prior to virus challenge. In experiments using CsH treatment, the media was replaced after 16 hrs treatment in order to avoid CsH toxicity^43^. At 48 hrs post-challenge with 2-part HIV-1 vectors, or at 72 hrs post-challenge with 3-part lentiviral vectors, cells were harvested for flow cytometric analysis, by pipetting (CD4^+^ T cells) or scraping (DCs and macrophages). Cells were pelleted at 500 × g for 5 min, and fixed in a 1:4 dilution of BD Cytofix Fixation Buffer with phosphate-buffered saline (PBS) without Ca^2+^ and Mg^2+^, supplemented with 2% FBS and 0.1% NaN_3_.

### Flow Cytometry

Data was collected on an Accuri C6 (BD Biosciences, San Jose, CA) and plotted with FlowJo software. Infectivity at each dilution, in each condition (CA mutant, CypA inhibitor, or CypA knockdown) was compared to the infectivity of WT CA in the control condition. Dilutions yielding infectivity greater than 30% GFP+ cells were excluded from analysis on the assumption that these were out of the linear range, according to the Poisson distribution.

### Statistical analysis

Experimental n values and information regarding the statistical tests can be found in the figure legends. The data of infectivity assay using single-cycle viruses including at least three independent donors were statistically analyzed by using two-tailed paired t-test compared to the control condition or the indicated condition for each donor. The data of PLA quantification were assessed for statistical significance using two-tailed unpaired t-test to compare two conditions as indicated in Fig. 4. All statistical analyses were performed using PRISM 7.0 (GraphPad Software, La Jolla, CA).

### Quantitative PCR for viral late reverse transcriptase product

Total DNA was extracted from cells using DNeasy Blood & Tissue Kit (Qiagen, Hilden, Germany), following the manufacturer’s instruction. Late RT products were detected with TaqMan system using the primers pWPTS J1B fwd and pWPTS J2 rev with the late RT probe (LRT-P)^44^. Mitochondrial DNA was used for normalization with the following primer/probe set: MH533, MH534 and Mito probe^45^. The primer and probe sequences are specified in Supplementary Table 3. The quantitative PCR was performed in 20 μL reaction mix containing 1× TaqMan Gene Expression Master Mix (Applied Biosystems), 900 nM each primer, 250 nM TaqMan probe and 30 to 50 ng template DNA. After an initial incubation at 50 °C for 2 min and the second incubation at 95 °C for 10 min, 45 cycles of amplification were carried out at 95 °C for 15 sec followed by 1 min and 30 sec at 60 °C. Real-Time PCR reactions were run on a CFX96^™^ thermal cycler (Bio-Rad).

### qRT-PCR

Total RNA was isolated in TRIzol reagent followed by RNA purification with RNeasy Plus Mini kit (Qiagen). First-strand cDNA was generated using SuperScript^™^ VILO^™^ Master Mix (Thermo Fisher) with random hexamers, in accordance with manufacturer’s instructions. Duplex qPCR was performed in 20 μL reaction mix containing 1× TaqMan Gene Expression Master Mix, 1× TaqMan Gene Expression Assay detecting TRIM5 (FAM dye-labeled, TaqMan probe ID #Hs01552559_m1), 1× TaqMan Gene Expression Assay targeting a housekeeping gene OAZ1 (VIC dye-labeled, primer-limited, TaqMan probe ID #Hs00427923_m1). Amplification was on a Biorad CFX96 real-time PCR detection system, using: 95 °C for 10 min, then 45 cycles of 95 °C for 15 sec and 60 °C for 60 sec.

### Western blot

Cells were lysed in Hypotonic Lysis Buffer: 20 mM Tris-HCl, pH 7.5, 150 mM NaCl, 10 mM EDTA, 0.5% NP-40, 0.1% Triton X-100, and cOmplete mini protease inhibitor (Sigma-Aldrich) for 20 min on ice. The lysates were mixed 1:1 with 2 × Laemmli buffer containing 1:20-diluted 2-mercaptoethanol, boiled for 10 min, and centrifuged at 16,000 × g for 5 min at 4°C. Samples were run on 4–20% SDS-PAGE and transferred to nitrocellulose membranes. Membrane blocking, as well as antibody binding were in TBS Odyssey Blocking Buffer (Li-Cor, Lincoln, NE). Primary antibodies used were rabbit anti-CypA (1:10,000 dilution; Enzo Life Sciences, Farmingdale, NY, catalogue #BML-SA296) and mouse anti-ß-actin (1:1,000 dilution; Abcam, Cambridge, UK, catalogue #ab3280). Goat anti-mouse-680 (Li-Cor, catalogue #925-68070) and goat anti-rabbit-800 (Li-Cor, catalogue #925-32211) as secondary antibodies were used at 1:10,000 dilutions. Blots were scanned on the Li-Cor Odyssey CLx.

### Proximity ligation assay (PLA)

2.5 × 10^5^ macrophages were plated on 12 mm coverslips (Warner Instrument, Hamden, CT, catalogue #CS-12R15) in 24-well plates. Cells were spinoculated at 1,200 × g using 6 × 10^8^ RT unit/mL of 3-part lentiviral vector (pALPS-GFP, p8.9NΔSB, and pMD2.G, generated as above) at 13°C for 2 hrs. Media was replaced with RPMI-HS complete containing 2 μM MG132, and either DMSO or 5 μM CsA, and cells were incubated at 37° for 2 hrs. Coverslips were fixed with 3.7% formaldehyde (ThermoFisher) in 0.1 M PIPES, pH 6.8, for 5 min at room temperature, and then incubated at room temperature for 1 hr in PBS containing 0.1% saponin, 10% donkey serum, 0.01% sodium azide, mouse anti-TRIM5α antibody (NIH AIDS Reagent Program, catalogue #12271) at a 1:750 dilution and rabbit anti-HIV-1 CA (p24) antibody (Abcam, catalogue #ab32352) at a 1:400 dilution. The samples were processed further using a Duolink^®^ In Situ Red kit (Sigma-Aldrich), following the instructions of the manufacturer. Then samples were incubated with 10 μM phalloidin (FITC) (Enzo Life Science) and 1 mg/mL Hoechst 33342 (Invitrogen), in PBS containing 10% donkey serum and 0.01% sodium azide for 30 min at room temperature. Coverslips were mounted on slides and stored at −20 °C. Interaction was detected as fluorescent spots (λ_excitation/emission_ 598/634 nm). λ_excitation/emission_ 475/523 nm and λ_excitation/emission_ 390/435 nm were used to detect phalloidin and Hoechst, respectively. Z-stack images were collected with a DeltaVision wide-field fluorescent microscope (Applied Precision, GE) and deconvolved with SoftWoRx deconvolution software (Applied Precision, GE). All images were acquired under identical acquisition conditions and analyzed by Imaris 8.3.1 (Bitplane). Three dimensional representations were constructed by using the Easy 3D function (Imaris 8.3.1).

### Challenge with replication-competent HIV-1

5 × 10^5^ macrophages per well in 12-well plates were challenged with 10^8^ RT units of HIV-1 for 2 hrs, in the presence of CsA or DMSO solvent, as indicated. Macrophage experiments used NL4-3_MAC_; pNL4-3, in which *env* was replaced from the end of the signal peptide to the *env* stop codon, with macrophage-tropic *env* from GenBank #U63632.1. 3 days after secondary stimulation with CD3/CD28, 10^6^ CD4^+^ T cells per well in 48-well plates were challenged with 2 × 10^7^ RT units of HIV-1 for 2 hrs, in the presence of GS-CypAi3 or DMSO solvent, as indicated. CD4^+^ T cell experiments used HIV-1_ZM249M_, a clade C transmission-founder strain. After HIV-1 challenge, cells were washed with fresh media and resuspended in 1 mL of RPMI-HS complete for macrophages or RPMI-FBS complete containing 50 U/mL hIL-2 for CD4^+^ T cells. Where indicated, culture media also contained CsA, GS-CypAi3, or DMSO solvent. Every 2-3 days, culture supernatant was harvested to measure RT activity.

Supplementary information is available in the online version of the paper.

## Acknowledgements

We thank Tomas Cihlar, Beatrice Hahn, Stephane Hausmann, Eric Hunter, Richard Mackman, Massimo Pizzato, and Stephen Yant for reagents. We are also grateful to anonymous blood donors who contributed leukocytes to this study. This work was supported by NIH Grants 5R01AI111809, 5DP1DA034990, and 1R01AI117839, to J.L.

## Author Contributions

K.K. and J.L. designed the experiments. K.K., A.D., S.K., L.Y., W.E.D., S.M.M., and C.C. conducted the experiments. C.S. and E.M.C provided advice and technical expertise. All authors analyzed the data. K.K. and J.L. wrote the manuscript.

## Author information

The authors declare no competing financial interests. Correspondence and requests for materials should be addressed to J.L.

## Data availability statement

The plasmids described in Supplementary Table 1 are available at www.addgene.com. All data generated or analyzed during this study are presented in the paper or in the Supplementary Information file, as well as are available from the corresponding author upon reasonable request.

