## Supplementary material for "Cyclophilin A protects HIV-1 from restriction by human TRIM5α": Supp Figs 1-4 Supp Tables 1-3 (from another version of this preprint: version 1, 2019-03-25)

Kyusik Kim, Ann Dauphin, Sevnur Komurlu, Leonid Yurkovetskiy, William E. Diehl, Sean M. McCauley, Claudia Carbone, Anastasia Selyutina, Angel Bulnes-Ramos, Felipe Diaz-Griffero, Caterina Strambio-De-Castillia, Edward M. Campbell, and Jeremy Luban

**Including:**

**Extended Data Figures 1, 2, 3 and 4 with corresponding Figure Legends.**

**Supplementary Tables 1, 2 and 3**

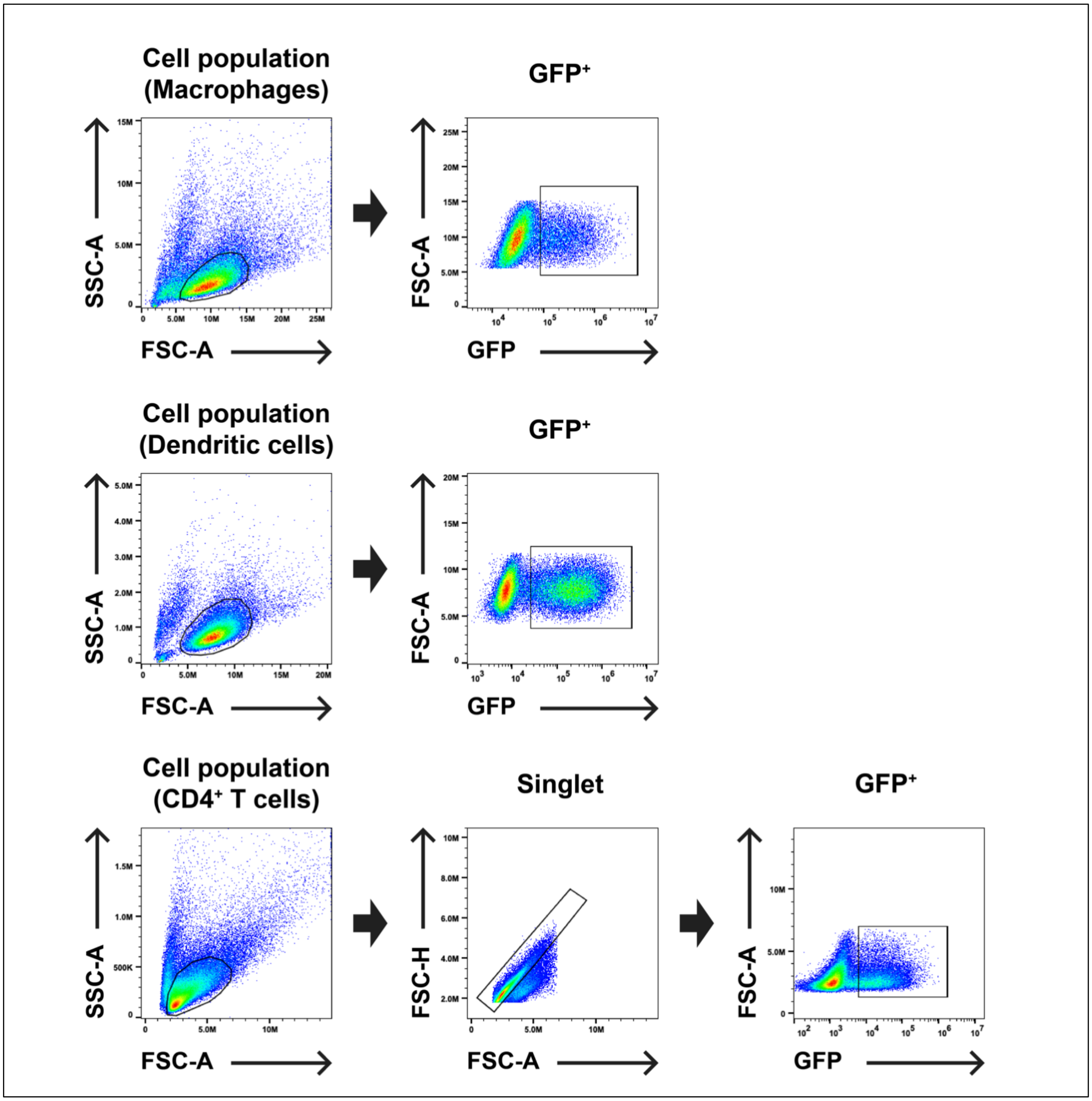


**Extended Data Figure 1. Gating strategy for flow cytometry experiments assessing single cycle infectivity.** Macrophage and dendritic cell populations, previously enriched as per the methods, were gated by SSC-A vs. FSC-A, as indicated, and then the $GFP^+$ population was plotted vs. FSC-A. Enriched $CD4^+$ T cells were gated by SSC-A vs FSC-A, as indicated, then a singlet population was gated from FSC-H vs. FSC-A, and finally $GFP^+$ cells were plotted vs. FSC-A.

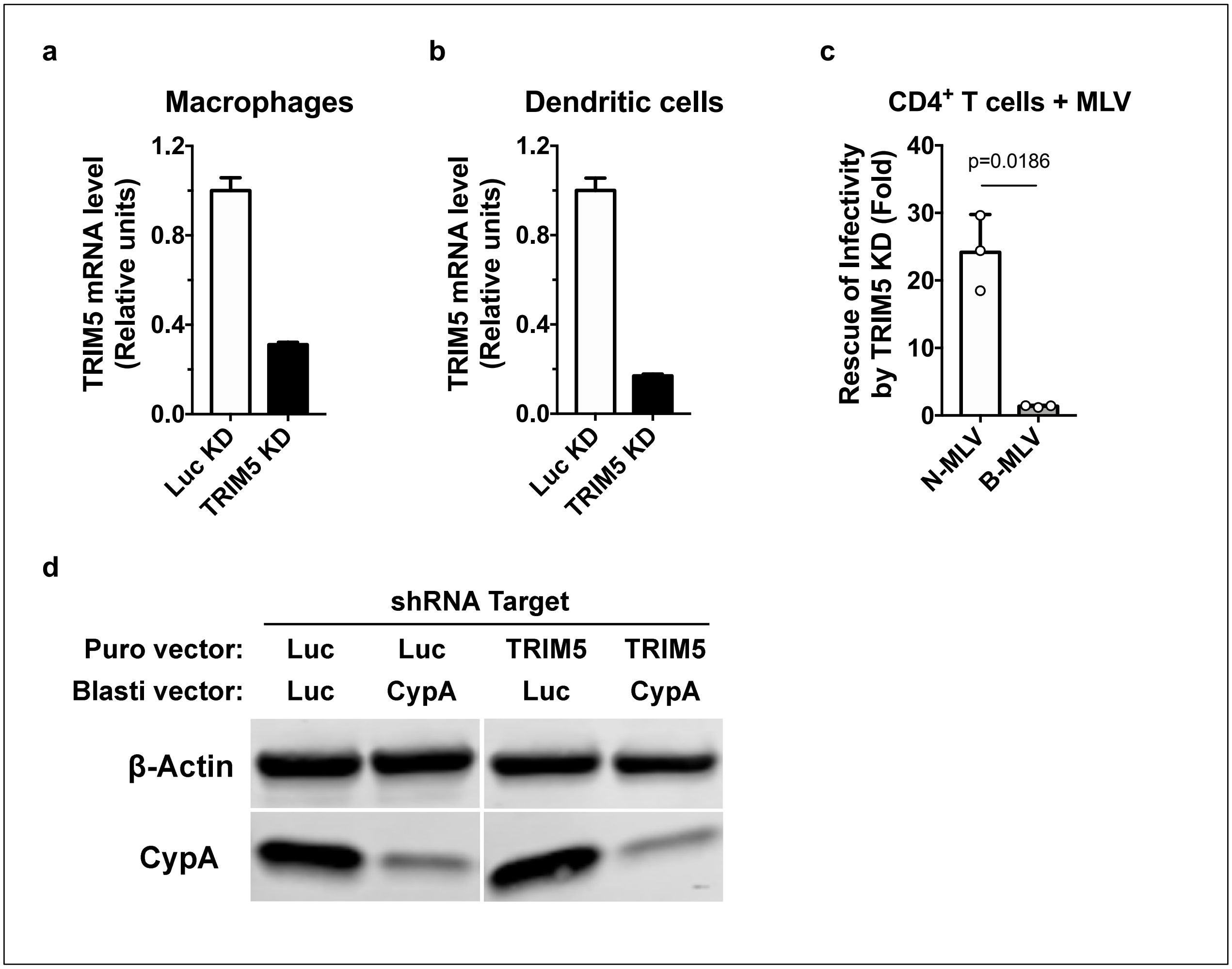


**Extended Data Figure 2. Assessment of shRNA-mediated knockdown in primary human blood cells. a**-**c**, Lentiviral vectors containing puromycin N-acetyltransferase (PAC) and shRNA targeting TRIM5 or Luc were used to transduce macrophages (**a**), dendritic cells (**b**), or $CD4^+$ T cells (**c**). At 3 days post-transduction, cells were selected with puromycin for 3 days. Total RNA was isolated from the macrophages and dendritic cells, followed by cDNA synthesis, and qPCR with TaqMan detection of TRIM5 and the housekeeping gene OAZ1, for normalization (**a** and **b**). The selected $CD4^+$ T cells were challenged with N- or B-MLV vector harboring GFP reporter for 3 days. Flow cytometry was used to assess the percentage of $GFP^+$ cells. The infectivity of each vector in TRIM5 knockdown cells was normalized to the Luc control condition (**c**). **d**, Macrophages were simultaneously transduced with two lentiviral vectors, the first expressing shRNA targeting TRIM5 or Luc with PAC, and the second expressing shRNA targeting CypA or Luc with blasticidin S-deaminase (bsd). After selection with both antibiotics, CypA and β-actin proteins were detected by western blot.

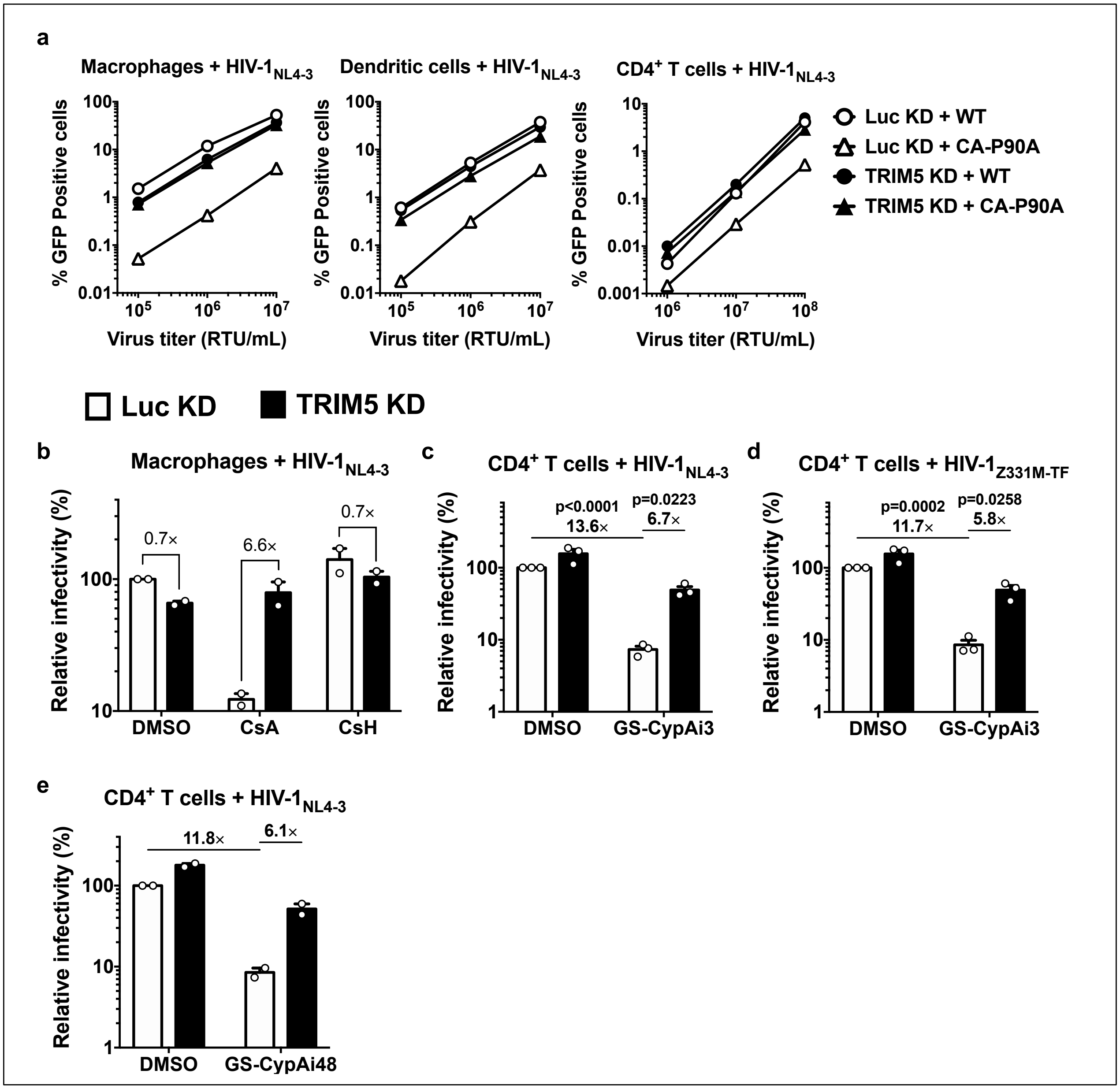


**Extended Data Figure 3. CA-CypA interaction promotes HIV-1 transduction by inhibiting TRIM5 activity in primary human blood cells.** **a**, Raw infectivity data for single cycle viruses, prior to normalization of infectivity to control condition. **b**, Macrophages expressing shRNA targeting TRIM5 or Luc were challenged with single-cycle, VSV G-pseudotyped, HIV-$1_{NL4-3}$GFP in the presence of 8 μM CsA, 8 μM CsH, or DMSO solvent (mean ± SEM, n = 2 donors). **c** and **d**, TRIM5 or Luc knockdown $CD4^+$ T cells were challenged with single-cycle HIV-$1_{NL4-3}$GFP (**c**) or HIV-$1_{Z331M-TF}$GFP (**d**) in the presence of 2.5 μM GS-CypAi3 or DMSO solvent alone (mean ± SEM, n = 3 donors for each). **e**, HIV-$1_{NL4-3}$GFP was used to challenge TRIM5 or Luc knockdown $CD4^+$ T cells with 2.5 μM GS-CypAi48 or DMSO solvent alone (mean ± SEM, n = 2 donors). Flow cytometry was used to measure the percentage of $GFP^+$ cells, followed by normalization to WT in Luc knockdown cells. Significance was determined by two-tailed, paired t-test for data generated from at least three donors.

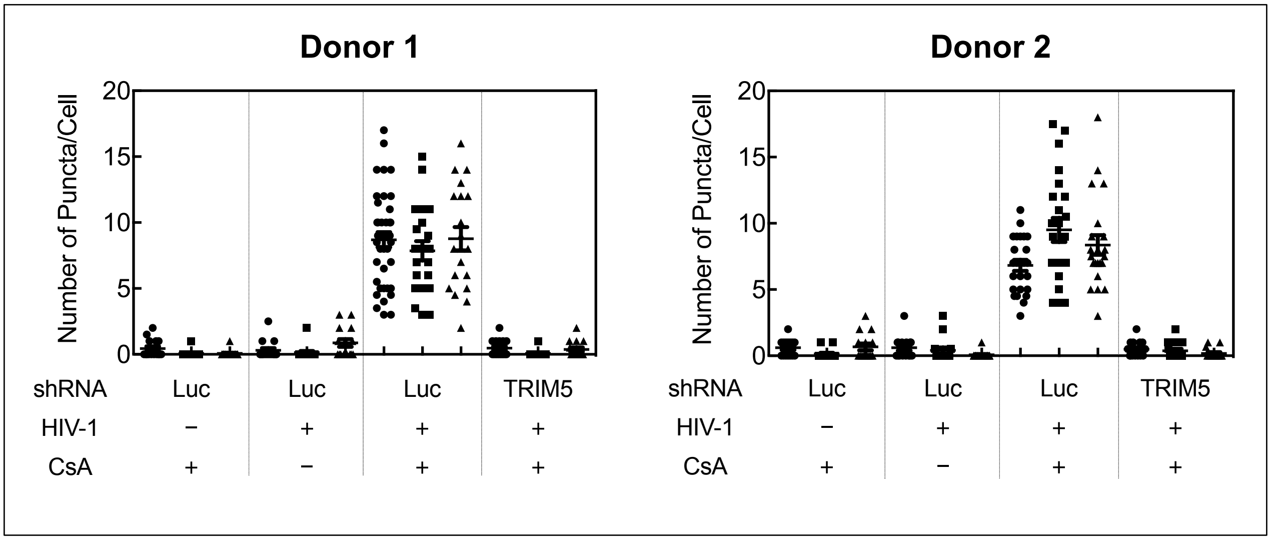


**Extended Data Figure 4. Association of endogenous TRIM5α with HIV-1 CA in primary human macrophages.** TRIM5 or Luc knockdown macrophages were challenged with VSV G-pseudotyped, HIV-$1_{NL4-3}$GFP in the presence of 5 μM CsA or DMSO solvent for 2 hrs. Proximity ligation assay (PLA) was then performed using anti-CA (p24) and anti-TRIM5α antibodies. Shown are the number of PLA puncta per cell, in each of three technical replicates, under each of the indicated conditions, for cells from two blood donors.

**Supplementary Table 1. Plasmids used in this study.**

| Plasmid Name | Purpose | Notes | Addgene # |
|---|---|---|---|
| pAIP-hGMCSF-co | Stable 293 cell line expressing human cytokine GM-CSF | SFFV promoter expresses codon-optimized human GM-CSF and puromycin N-acetyltransferase | 74168 |
| pAIP-hIL4-co | Stable 293 cell line expressing human cytokine IL4 | SFFV promoter expresses codon-optimized human IL4 and puromycin N-acetyltransferase | 74169 |
| pSIV-∆psi/∆env | $SIV_{MAC251}$ *gag-pol/vpx* | Production of SIV-VLPs containing Vpx protein | Pending |
| pMD2.G | VSV Glycoprotein | Pseudotype HIV-1 vectors with VSV Glycoprotein | 12259 |
| psPAX2 | HIV-1 *gag-pol* | Encodes gag structural proteins and pol enzymes to generate virion particles to generate 3-part lentiviral vector | 12260 |
| p8.9 N∆SB | HIV-1 *gag-pol* | Minimal HIV-1 packaging plasmid for gag and pol expression | Pending |
| p8.9 N∆SB P90A | HIV-1 *gag-pol* | p8.9 N∆SB bearing P90A mutant capsid sequence | Pending |
| p8.9 N∆SB D185K / D186L | HIV-1 *gag-pol* | p8.9 N∆SB bearing RT-D185K/D186L mutant reverse transcriptase sequence | Pending |
| pALPS GFP | GFP reporter of lentiviral vector | SFFV promoter expresses GFP | 101323 |
| pWPTS GFP | Lentivector for quantification of HIV-1 cDNA | Contains a unique loxP sequence within a region of the 3' LTR U3. EF1α promoter expresses GFP | 12255 |
| pAPM-CoE-D4-miR30-L1221 | Lentivector for luciferase (control) knockdown | SFFV promoter expresses puromycin N-acetyltransferase and miR30-based shRNA targeting luciferase.<br>Target sequence: 5′-TACAAACGCTCTCATCGACAAG-3′; negative control for knockdown | 115846 |
| pAPM-CoE-D4-miR30-TRIM5 | Lentivector for TRIM5 knockdown | SFFV promoter expresses puromycin N-acetyltransferase and miR30-based shRNA targeting TRIM5.<br>Target sequence: 5′-TGCCAAGCATGCCTCACTGCAA-3′ | Pending |
| pABM-CoE-D4-miR30-L1221 | Lentivector for luciferase (control) knockdown | SFFV promoter expresses blasticidin S-deaminase and miR30-based shRNA targeting luciferase.<br>Target sequence: 5′-TACAAACGCTCTCATCGACAAG-3′; negative control for knockdown | Pending |
| pABM-CoE-D4-miR30- | Lentivector for CypA knockdown | SFFV promoter expresses blasticidin S-deaminase and miR30-based shRNA | Pending |

| | | | |
|---|---|---|---|
| CypA | | targeting cyclophilin A.<br>Target sequence: 5′-CTGGATTGCAGAGTTAAGTTTA-3′ | |
| pPU-Empty-miR30-L1221 | All-in-one knockdown-rescue lentivector for luciferase (control) knockdown | SFFV promoter expresses puromycin N-acetyltransferase and miR30-based shRNA targeting luciferase.<br>Target sequence: 5′-TACAAACGCTCTCATCGACAAG-3′; negative control in knockdown-rescue experiment | Pending |
| pPU-Empty-miR30-TRIM5 | All-in-one knockdown-rescue lentivector for TRIM5 knockdown | SFFV promoter expresses puromycin N-acetyltransferase and miR30-based shRNA targeting TRIM5.<br>Target sequence: 5′-TGCCAAGCATGCCTCACTGCAA-3′ | Pending |
| pPU-huTRIM5α-miR30-L1221 | All-in-one knockdown-rescue lentivector for huTRIM5α expression and luciferase (control) knockdown | SFFV promoter expresses puromycin N-acetyltransferase, exogenous huTRIM5α protein and miR30-based shRNA targeting luciferase.<br>Target sequence: 5′-TACAAACGCTCTCATCGACAAG-3′ | Pending |
| pPU-huTRIM5α-miR30-TRIM5 | All-in-one knockdown-rescue lentivector for huTRIM5α expression and TRIM5 knockdown | SFFV promoter expresses puromycin N-acetyltransferase, exogenous huTRIM5α protein and miR30-based shRNA targeting TRIM5.<br>Target sequence: 5′-TGCCAAGCATGCCTCACTGCAA-3′ | Pending |
| pNL4-3/Mac-tropic Env | Full-length infectious HIV-$1_{MAC}$ | Encodes HIV-$1_{NL4-3}$ proviral sequences containing macrophage-tropic JR-FL *env* | Pending |
| pLXIN GFP | GFP reporter of MLV-based vector | MLV-based transfer vector expressing GFP and neomycin resistance protein | Pending |
| pCIG3 N | N-tropic MLV *gag-pol* | MLV packaging plasmid for N-tropic MLV gag-pol expression | Pending |
| pCIG3 B | B-tropic MLV *gag-pol* | MLV packaging plasmid for B-tropic MLV gag-pol expression | Pending |
| pUC57mini ZM249M | HIV-1 clade C molecular clone | Molecular clone of transmitted/founder virus HIV-$1_{ZM249M}$ | Pending |
| pUC57mini ZM249M P90A | HIV-1 clade C molecular clone | Molecular clone of transmitted/founder virus HIV-$1_{ZM249M}$ bearing P90A capsid mutation | Pending |
| pUC57mini Z331M-TF | HIV-1 clade C molecular clone | Molecular clone of transmitted/founder virus HIV-$1_{Z331M-TF}$ | Pending |

| | | | |
|---|---|---|---|
| pUC57mini ZM249M Δenv eGFP | HIV-1 clade C molecular clone | Molecular clone of transmitted/founder virus HIV-$1_{ZM249M}$ with deletion of 79 nucleotides following the env signal peptide and GFP in place of nef | Pending |
| pUC57mini ZM249M P90A Δenv eGFP | HIV-1 clade C molecular clone | Molecular clone of transmitted/founder virus HIV-$1_{ZM249M}$ containing P90A capsid mutation, with deletion of 79 nucleotides following the env signal peptide and GFP in place of nef | Pending |
| pUC57mini Z331M-TF Δenv eGFP | HIV-1 clade C molecular clone | Molecular clone of transmitted/founder virus HIV-$1_{Z331M\text{-}TF}$ with deletion of 79 nucleotides following the env signal peptide and GFP in place of nef | Pending |

**Supplementary Table 2. Drugs and reagents.**

| Reagents | Source | Identifier |
|---|---|---|
| DMEM, high glucose | ThermoFisher | 11995073 |
| Fetal Bovine Serum | GE Healthcare Life Sciences | SH30087.03 |
| RPMI-1640 | ThermoFisher | 11875119 |
| Human AB+ Serum | Omega Scientific | HS-20 |
| HEPES | Corning | 25-060-CI |
| MEM Non-Essential Amino Acids | Corning | 25-025-CI |
| Sodium Pyruvate | Corning | 25-000-CI |
| GlutaMAX™-I | ThermoFisher | 35050-061 |
| Opti-MEM | ThermoFisher | 31985-062 |
| TransIT-LT1 | Mirus Bio | MIR2306 |
| BD Cytofix Fixation Buffer | BD Bioscience | 554655 |
| cOmplete mini protease inhibitor | Roche | 11836170001 |
| 2-mercaptoethanol | Invitrogen | 21985-023 |
| Laemmli Buffer | BioRad | 161-0737 |
| 4-20% gradient SDS-PAGE gels, 12 well | BioRad | 456-1095 |
| 0.45 mM nitrocellulose membrane | BioRad | 162-0112 |
| TBS Odyssey Blocking Buffer | Li-Cor | 927-50000 |
| Pierce™ 16% Formaldehyde (w/v), Methanol-free | ThermoFisher | 28906 |
| PIPES | Sigma-Aldrich | P1851 |
| Phalloidin (FITC) | Enzo Life Science | ALX-350-268-MC01 |
| Hoeschst 33342 | Invitrogen | H3570 |
| Cyclosporine A | Sigma-Aldrich | 30024 |
| Cyclosporine H | Sigma-Aldrich | SML1575 |
| GS-CypAi3 | Gilead | Non-commercial |
| GS-CypAi48 | Gilead | Non-commercial |
| MG132 | Sigma-Aldrich | M7449 |

**Supplementary Table 3. qPCR primers and probes for Late RT quantification**

| Primer or Probe name | Sequence | Note |
|---|---|---|
| pWPT J1B Fwd | 5’-GCATACATTATACGAAGTTATGCTGC-3’ | Primer to detect late RT product |
| pWPT J2 Rev | 5’-GCCGTGCGCGCTTCAGCAAGC-3’ | Primer to detect late RT product |
| LRT-P (Probe) | 5’-(FAM)-CAGTGGCGCCCGAACAGGGA-(TAMRA)-3’ | Probe to detect late RT product |
| MH533 | 5′-ACCCACTCCCTCTTAGCCAATATT-3′ | Primer to detect mitochondrial DNA |
| MH534 | 5′-GTAGGGCTAGGCCCACCG-3′ | Primer to detect mitochondrial DNA |
| Mito-P (Probe) | 5′-(FAM) CTAGTCTTTGCCGCCTGCGAAGCA (TAMRA)-3′ | Probe to detect mitochondrial DNA |
